# Rare and Common Variants Associated with Alcohol Consumption Identify a Conserved Molecular Network

**DOI:** 10.1101/2024.02.26.582195

**Authors:** Brittany S. Leger, John J. Meredith, Trey Ideker, Sandra Sanchez-Roige, Abraham A. Palmer

## Abstract

Genome-wide association studies (GWAS) have identified hundreds of common variants associated with alcohol consumption. In contrast, rare variants have only begun to be studied for their role in alcohol consumption. No studies have examined whether common and rare variants implicate the same genes and molecular networks. To address this knowledge gap, we used publicly available alcohol consumption GWAS summary statistics (GSCAN, N=666,978) and whole exome sequencing data (Genebass, N=393,099) to identify a set of common and rare variants for alcohol consumption. Gene-based analysis of each dataset have implicated 294 (common variants) and 35 (rare variants) genes, including ethanol metabolizing genes *ADH1B* and *ADH1C*, which were identified by both analyses, and *ANKRD12, GIGYF1, KIF21B*, and *STK31*, which were identified only by rare variant analysis, but have been associated with related psychiatric traits. We then used a network colocalization procedure to propagate the common and rare gene sets onto a shared molecular network, revealing significant overlap. The shared network identified gene families that function in alcohol metabolism, including *ADH, ALDH, CYP*, and *UGT*. 74 of the genes in the network were previously implicated in comorbid psychiatric or substance use disorders, but had not previously been identified for alcohol-related behaviors, including *EXOC2, EPM2A, CACNB3*, and *CACNG4*. Differential gene expression analysis showed enrichment in the liver and several brain regions supporting the role of network genes in alcohol consumption. Thus, genes implicated by common and rare variants identify shared functions relevant to alcohol consumption, which also underlie psychiatric traits and substance use disorders that are comorbid with alcohol use.

## Introduction

Alcohol use disorder (**AUD**) is a highly heritable (Verhulst, Neale and Kendler, 2015) disease with a heavy public health burden (MacKillop *et al*., 2022). AUD can be viewed as the endpoint of a series of transitions, which begin with the initiation of use, regular alcohol consumption that continues with the escalation to hazardous drinking, and culminates in compulsive harmful use that persists despite negative consequences (Sanchez-Roige, Palmer and Clarke, 2020). As such, alcohol consumption is frequently studied as a proxy for AUD, as it is a component of AUD, and is a quantitative trait that is widely measured, providing large sample sizes for genetic studies. In particular, genome-wide association studies (**GWAS**) have identified numerous common variants that contribute to alcohol consumption, AUD, and related traits (Clarke *et al*., 2017; Liu *et al*., 2019; Saunders *et al*., 2022).

Recently, GWAS of other psychiatric disorders have extended their reach to rare variants. Variants with deleterious effects are under negative selection (Gibson, 2012), thus rare variants are predicted to have higher penetrance and higher effect sizes in disease than common variants (Kimura *et al*., 2021). Rare variants also tend to have developed more recently than common variants, leading to fewer variants being in linkage disequilibrium with rare variants. This makes the association of rare variants with genes less ambiguous, and increases the interpretability of rare variants, compared with common variants (Sazonovs and Barrett, 2018). Rare variant studies have revealed that these variants influence risk for multiple psychiatric disorders, including intellectual disability, autism spectrum disorder, and schizophrenia (Ganna *et al*., 2018; Charney *et al*., 2019; Antaki *et al*., 2022; Singh *et al*., 2022; Fu *et al*., 2023; Weiner *et al*., 2023). Because they are uncommon, rare variants are best identified using sequencing in conjunction with large sample sizes (Manolio *et al*., 2009; Backman *et al*., 2021; Wang *et al*., 2021; Karczewski *et al*., 2022). Although a few exome sequencing studies and rare variant studies for alcohol phenotypes have been undertaken (Vrieze *et al*., 2014; Marees *et al*., 2018; Curtis, 2022; Ahangari *et al*., 2023) the contribution of rare variation on alcohol behaviors remains poorly characterized, as does the relationship between common and rare variants.

One way to address the relationship between common and rare variants is by using biological knowledge networks. These networks contain information about the molecular interactions among genes and their products, both broadly and in disease contexts (Jia and Zhao, 2014; Farris, Harris and Ponomarev, 2015; Fong *et al*., 2019; Rosenthal *et al*., 2023). While the interplay between rare and common variant-implicated genes has been studied in network space for other psychiatric traits (Gilman *et al*., 2011; Ben-David and Shifman, 2012; Chang *et al*., 2018), it has not been studied for alcohol-related traits or other substance use disorders (**SUD**). Based on evidence from comparisons of common and rare variants for other psychiatric traits, we hypothesized that the same genes and molecular pathways would be identified by both approaches.

To test this hypothesis, we assembled data from UK Biobank (**UKB**) pertaining to both common and rare variants that are associated with alcohol consumption. We then used a network approach to investigate the biological overlap between common and rare variants for alcohol consumption. This approach allowed us to compare their relative contributions at the variant, gene, and molecular pathway levels.

## Results

### Common and Rare Variant Associations

We obtained GWAS summary statistics from GSCAN, which recently performed a meta-analysis of alcohol consumption in Europeans (Saunders *et al*., 2022) (n=666,978, **Figure 1A**). 501 independent (r^2^) common (MAF>0.05) variants were significantly associated with alcohol consumption (*p*<5×10^−8^) (Saunders *et al*., 2022). Genome-wide significant rare variants were obtained from Genebass’s recent analysis of 393,099 individuals from the UKB (Karczewski *et al*., 2022) (**Figure 1B**). Three rare variants were significantly associated with alcohol consumption (*p*<8×10^−9^): one potential loss of function (pLoF) variant in *ADH1C*, a missense variant in *ADH1B*, and a synonymous mutation in *C4orf54* (**Table S1**). The mutations in *ADH1C* and *C4orf54* were both protective. *ADH1C* and *ADH1B* both have known roles in ethanol metabolism (Tolstrup *et al*., 2008; Le Daré, Lagente and Gicquel, 2019), but despite *C4orf54* being associated with substance use disorders in prior GWAS (Sollis *et al*., 2023), its function is poorly understood.

**Figure 1.**
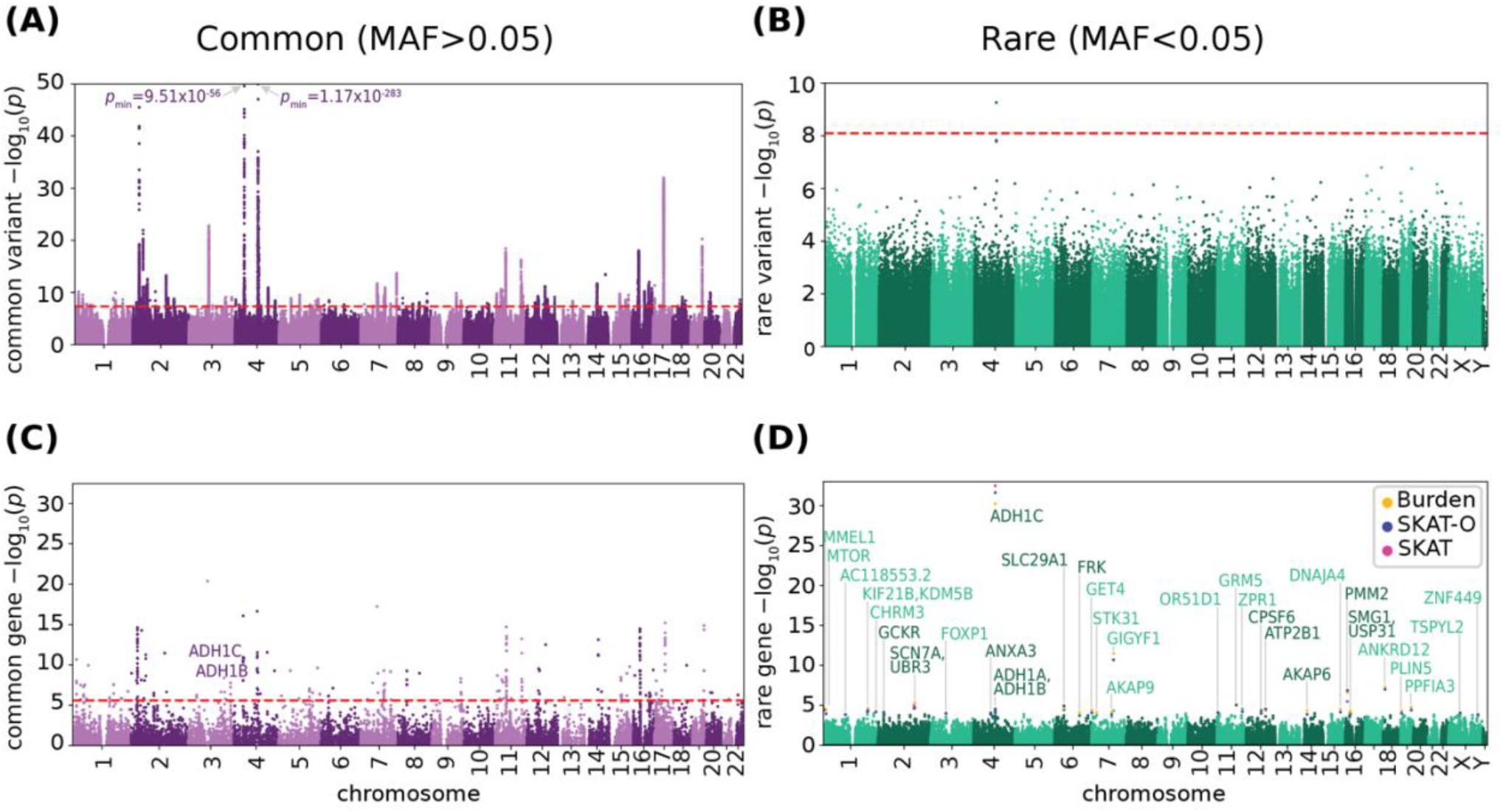
Common and rare variants mediate alcohol consumption. Manhattan plot of (**A**) common variants and (**B**) rare variants associated with alcohol consumption. Significance cutoff indicated in red (common: *p*<5×10^−8^; rare: *p*<8×10^−9^). *p*-value for peaks outside of range labeled. Rare variant MAC>2. (**C**) Manhattan plot of alcohol consumption common variant-implicated genes. Significance cutoff (*p*<2.6×10^−6^) indicated in red. Significant genes that overlap with rare-variant implicated genes are labeled. (**D**) Porcupine plot of genes calculated by burden test, SKAT-O, and SKAT algorithms from rare variants. Significantly associated genes (FDR<0.25) for each test are labeled and colored in yellow, blue, and pink, for burden, SKAT-O, and SKAT, respectively. See Figure S1a for individual manhattan plots for each test.

### Common and Rare Gene-Level Associations

Common loci were assigned to genes based on proximity using MAGMA (de Leeuw *et al*., 2015), identifying 294 genes (**Figure 1C**; **Table S2**, *p*<2.6×10^−6^). Rare variants were previously aggregated (Karczewski *et al*., 2022) into gene level associations using SKAT, SKAT-O, and a variant burden test (Karczewski *et al*., 2022). These tests identified four genes that were significantly correlated with alcohol consumption via both SKAT-O and burden tests (*p*_SKAT-O_<2.5×10^−7^; *p*_burden_<6.7×10^-7^): *ADH1C, PMM2, GIGYF1*, and *ANKRD12*. Only *ADH1C* was significantly associated by SKAT (*p*_SKAT_<2.5×10^−7^), and was the only gene previously associated with alcohol-related traits by common gene analysis.

We also considered a more lenient cutoff for genes from rare variants (FDR<0.25, **Figure 1D, Figure S1A, Table S3**), which identified 35 genes across all tests. 20 genes were identified by both SKAT-O and the burden test (**Figure S1B**), however, only *ADH1C* and *PMM2* were significant in all tests. 51% of genes were functionally annotated as loss of function, followed by missense and low confidence loss of function (40%), and the remaining 9% as synonymous (**Figure S1C**). 12 of these genes had previously been identified by common variants as mediating alcohol consumption and alcohol use traits in the GWAS catalog (Sollis *et al*., 2023) (**Table S3**; *p*=8.24×10^−33^, hypergeometric test). This includes alcohol dehydrogenase genes *ADH1A, ADH1B*, and *ADH1C*, and signaling genes *FOXP1, AKAP6, AKAP9, and GRM5*, highlighting the overlapping regulation of SUDs and psychiatric traits.

*ADH1B* and *ADH1C* were identified by both the rare and common gene-based analyses (**Figure 2A**, *p*=0.01, hypergeometric test).

**Figure 2.**
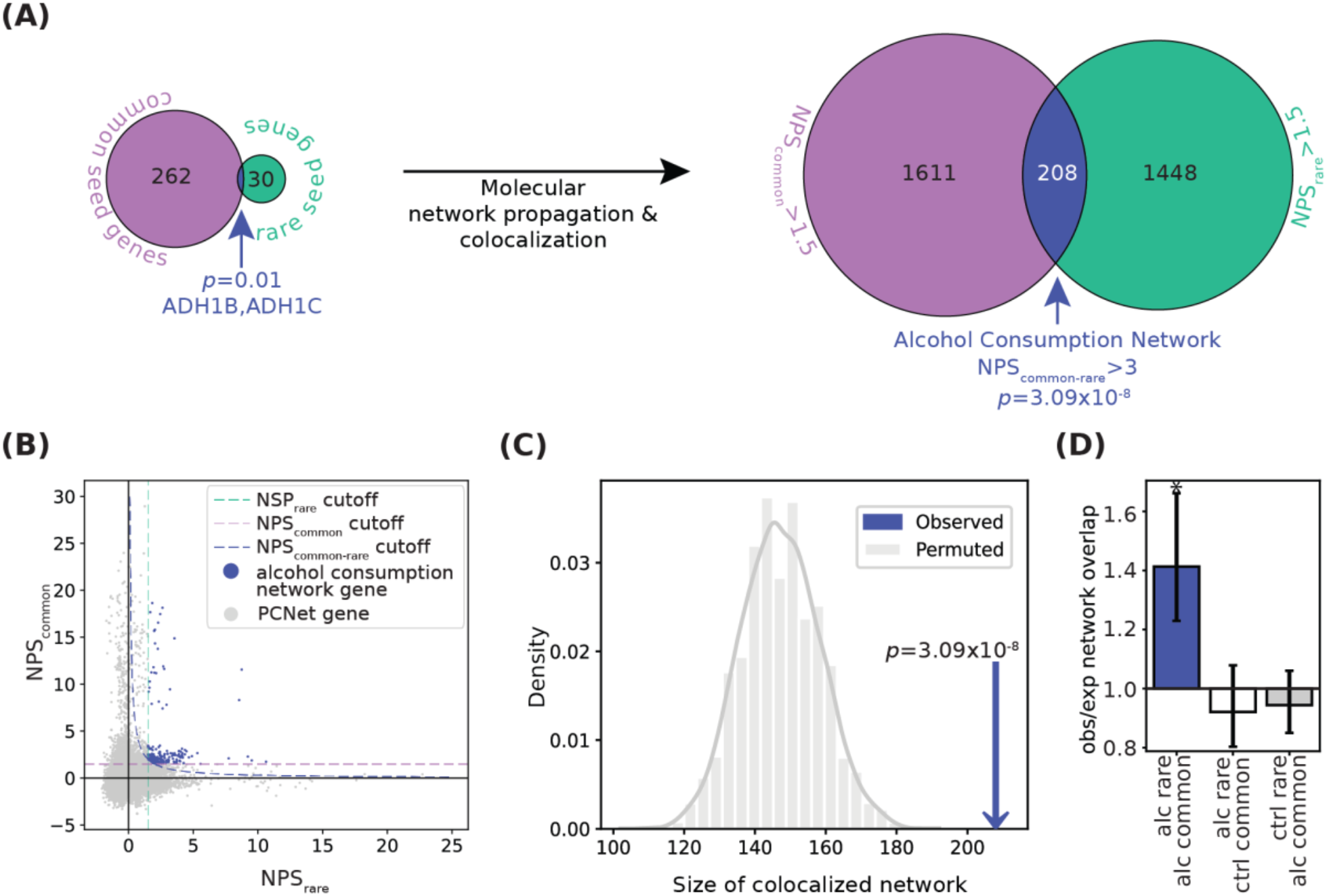
Convergence of rare and common variants on the network level. (**A**) Left, venn diagram showing overlap of common seed genes (purple) and rare seed genes (green). Overlapping genes are indicated in dark blue and labeled. Significance of overlap calculated via hypergeometric test. Right, Venn diagram of genes passing network proximity score (NPS) thresholds after network colocalization. Significance of intersection indicated calculated in C. (**B**) NPS_common_ and NPS_rare_ for all genes in PCNet, with genes passing all thresholds for the alcohol consumption network (NPS_common-rare_ > 3, NPS_common_ > 1.5, and NPS_rare_ > 1.5) shown in dark blue. Dotted lines indicate NPS thresholds. (**C**) Observed (dark blue arrow) versus expected (gray distribution) size of the Alcohol Consumption Network following 10,000 permutations of NPS labels. p-value calculated via Z-test. (**D**) The observed-to-expected ratio of colocalized network size for networks calculated from common and rare seed genes from alcohol consumption and from control trait FEV1 (forced expiratory volume per second). Vertical bars indicate 95% confidence intervals. Significance calculated by Z-test, Bonferroni corrected. * indicates *p*=3.09×10^−8^. See also Figure S2C and Table S5 for additional controls.

### Generation of the Alcohol Consumption Network

Next, we examined the molecular pathways wherein these alcohol consumption genes function (**Figure 2A**). We used the Parsimonious Composite Network (**PCNet**), a resource of 2.7 million pairwise associations among genes (Huang *et al*., 2018). PCNet is a consensus of 21 physical and functional interaction databases and integrates multiple lines of evidence, including protein-protein interactions, mRNA, protein co-expression across tissues, and literature curation.

We assigned network proximity scores (**NPS**) to each gene in PCNet using a random-walk algorithm, which computes the number of steps through the network to reach that gene from a set of seed genes. Seed genes from common and rare gene-set analyses were filtered for presence in PCNet, resulting in 264 common seed genes and 32 rare seed genes. NPS_common_ was calculated from common seed genes and NPS_rare_ was calculated from rare seed genes (**Figure S2A**). We then calculated the product of the two proximity scores to compute NPS_common-rare_ = NPS_common_ x NPS_rare_, and selected for high NPS_common_, NPS_rare_, and NPS_commonrare_ scores (**Figure 2B**). In this way, genes with the highest NPS_common-rare_ were close in the molecular network to both common and rare seed genes, even if they were not identified by the individual studies (**Table S4**).

We found that the alcohol consumption network contained significantly more genes (**Figure 2C**, *p*=3.09 x 10^−8^), and that the mean of NPS_common-rare_ was significantly higher than expected (**Figure S2C**, *p*=5.51 x 10^−6^). As a negative control, we produced networks using both the alcohol rare and common seed genes in conjunction with arbitrary traits; these negative controls did not produce networks that were larger than the permuted control (**Figure 2D, Table S5**). Additionally, when we considered a more stringent threshold for rare seed genes (*p*_SKAT-O_<2.5×10^−7^, n=4) we had similar results (**Figure S2C**). However, network colocalization was contingent upon *ADH1C*.

As shown in **Figure 3**, the alcohol consumption network contained 208 nodes, connected by 1,226 edges. 27 of 264 seed genes were maintained from the common seed genes. 5 of the 34 seed genes derived from rare variants were maintained into the network. *ADH1B* and *ADH1C*, which were seed genes for both common and rare, were both maintained into the network (**Table S4**).

**Figure 3.**
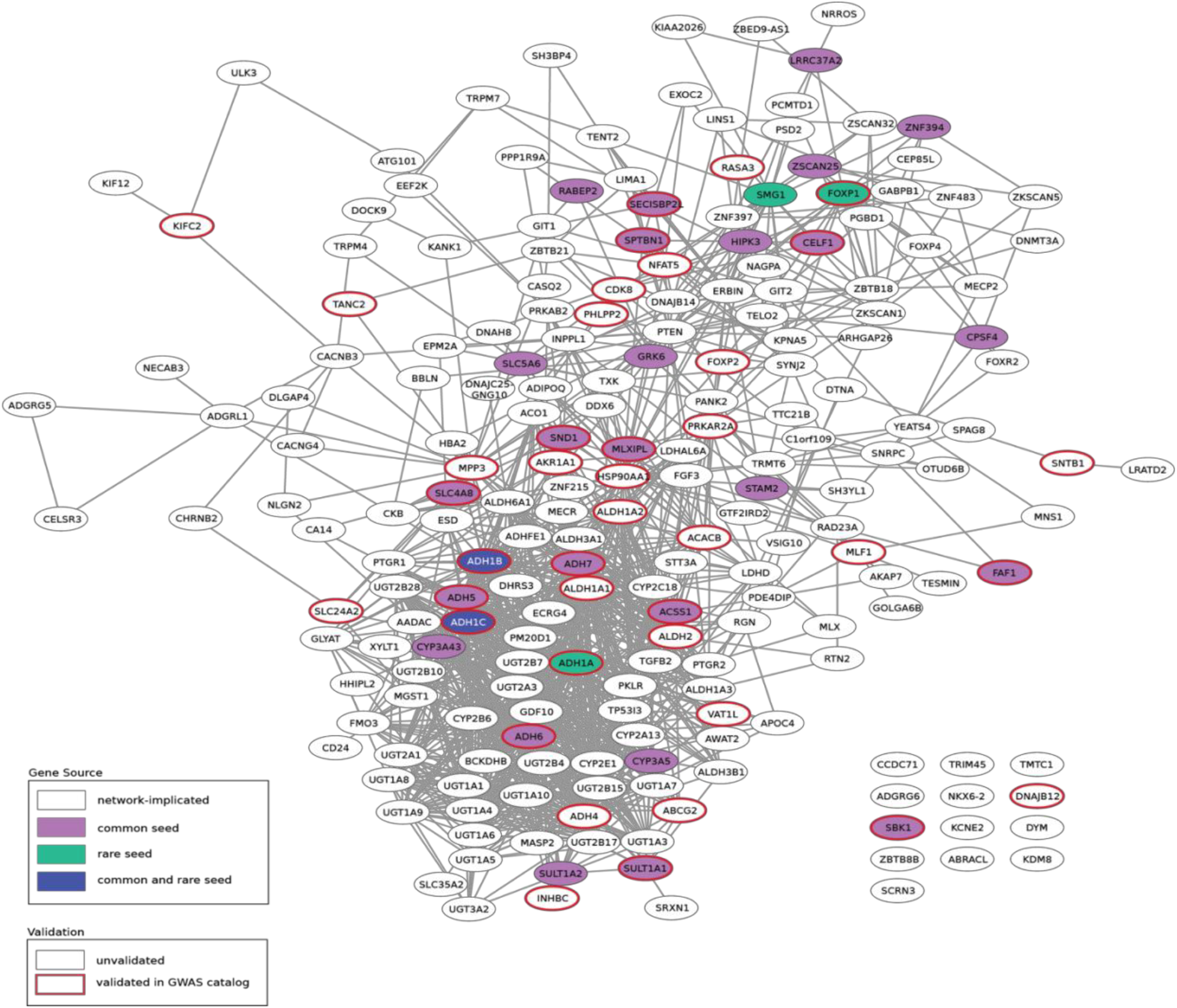
The Alcohol Consumption Network. Subnetwork of PCNet including all genes proximal to both rare and common alcohol consumption seed genes. Purple nodes indicate common seed genes, green nodes indicate rare seed genes, dark blue nodes indicate seeds in both sources, and white nodes are network-implicated genes. Edges maintained from PCNet. Red outlined nodes have previously been annotated in the GWAS catalog for alcohol use traits.

### The Structure of the Alcohol Consumption Network

One of the goals of generating the network shown in **Figure 3** is to identify the underlying biology identified by common and rare seed genes. Several gene families previously known to play a role in ethanol metabolism were present in the network (**Figure S3**). For example, 8 genes from the alcohol dehydrogenase (***ADH***) family (Le Daré, Lagente and Gicquel, 2019) and 7 aldehyde dehydrogenase (***ALDH***) family genes (Edenberg, 2007) are in the network. 6 cytochrome P450 (***CYP***) genes, which mediate about 10% of alcohol metabolism via the microsomal pathway (Hamitouche *et al*., 2006; Corella, 2012), were also in the network. In addition, genes from the non-oxidative ethanol metabolism pathways, which primarily function in phase II drug metabolism (Le Daré, Lagente and Gicquel, 2019), were also present. This includes 2 sulfotransferase (***SULT***) family genes, which metabolize ethanol into ethyl sulfate, and 18 genes in the UDP-Glycosyltransferase (***UGT***) superfamily, whose encoded proteins glucuronidate ethanol into ethyl glucuronide, a minor non-oxidative metabolite of ethanol (Walsham and Sherwood, 2014). Thus, the network recapitulates previously known biologies relevant to ethanol metabolism.

Another benefit of the network is the ability to identify relevant tissues. We found 25 tissues that were significantly enriched for differential gene expression (**Figure 4A, Table S3**), with high overlap of genes across tissues. Consistent with the presence of genes involved in ethanol metabolism in the network, the highest enrichment was in the liver and consisted of 115 genes, including 28 from the *ADH, ALDH, UGT, CYP*, and *SULT* families. In addition to the liver, numerous gastrointestinal tissues were also enriched: the gastrointestinal tract mediates absorption and gastric metabolism of alcohol, and chronic alcohol consumption may lead to inflammation and increased risk of gastrointestinal and esophageal cancers (Bode and Bode, 1997; Edenberg, 2007). As expected, all brain tissues were significantly enriched.

**Figure 4.**
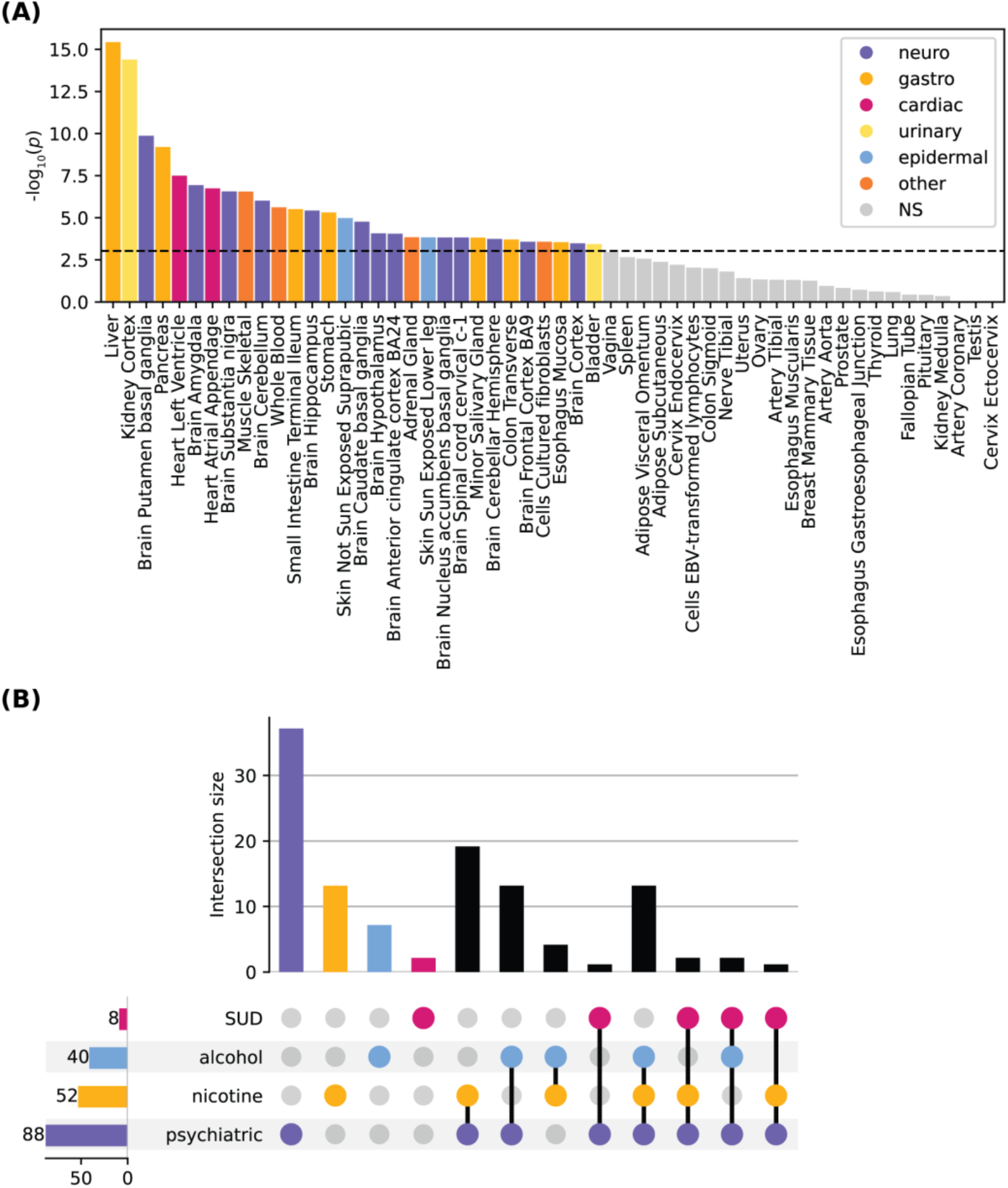
Validation of alcohol consumption network. (**A**) Enrichment of gene sets from the alcohol consumption network with bi-directional differential expression for 54 tissues from GTEX v8. Differentially expressed gene sets were defined by a two-sided t-tests per label, versus all remaining tissue types. Genes with *p* < 9.26×10^−4^ (Bonferroni corrected) and absolute log fold change ≥ 0.58 are selected as differentially expressed. Significance was calculated as the probability of the hypergeometric test. Tissues are colored by type, non-significant (NS) associations are indicated in gray. (**B**) Upset plot showing the overlap of genes in the alcohol consumption network that have previously been annotated in the GWAS catalog for alcohol use traits, nicotine use and smoking traits, other substance use disorders (SUD), and psychiatric traits.

To determine whether the genes had been previously implicated by common variants in alcohol use, other SUDs, and related psychiatric disorders, we examined annotations from the GWAS catalog (Sollis *et al*., 2023). Specifically, we considered annotations for alcohol use, smoking and nicotine use, and other SUDs, including opioid, cannabis, and polysubstance use, and related psychiatric disorders (**Figure 4B**). 201 of the 208 genes in the network are annotated in the GWAS catalog. Of these, 40 have been previously associated in alcohol use (*p*=1.56 x 10^−3^, hypergeometric test) and 52 network genes in smoking traits (*p*=0.046, hypergeometric test) (Karlsson Linnér *et al*., 2021). 8 were identified for SUDs and 88 for psychiatric traits, though the enrichment was not significant for either. Of the genes associated with these traits, many had annotations in multiple categories, such as *EPM2A, EXOC2, NFAT5*, and *SNTB1*. These findings highlight the neuropsychiatric function of the network and point to a shared underlying mechanism across alcohol and polysubstance use.

Finally, to determine whether these genes had been previously implicated by rare variants in alcohol use, we examined gene-level annotations from GeneBass of genes in the network (Karczewski *et al*., 2022). 6 of the 208 network genes (*ADH1C, AKAP7, ATG101, DTNA, NKX6-2*, and *SYNJ2*) were associated with secondary alcohol use traits by rare variants, including use status and frequency of use, negative societal impacts from use, and alcoholic liver disease. Only *ADH1C* was also associated with alcohol use traits by common variants. Notably, these genes, excluding *ATG101*, were all associated with other SUDs and psychiatric traits through common variants.

## Discussion

The contribution of common variants in mediating alcohol consumption has been well documented, while rare variants represent a new frontier that has recently become feasible due to the availability of large scale sequencing data. Prior rare variant analysis identified 4 genes at a stringent (*p*_SKAT-O_<2.5×10^−7^) and 35 genes at a lenient threshold (FDR<0.25), demonstrating the importance of rare variants for alcohol-related behaviors (**Figure 1**). We combined the findings from common and rare variants to determine whether they identify convergent biological networks (**Figure 2**). We identified a highly significant network (**Figure 3**). The network emphasized the role of ethanol metabolism, which was further supported by the tissue specific enrichment in both brain and liver (**Figure 4**), consistent with decades of research on the genetics of alcohol consumption.

The role of common variants in ethanol metabolizing enzymes is well established for alcohol consumption and related traits (Sanchez-Roige, Palmer and Clarke, 2020; MacKillop *et al*., 2022). Similarly, rare variant analysis of alcohol consumption identified *ADH1A, ADH1B*, and *ADH1C*, which have well documented roles in ethanol metabolism (Edenberg, 2007). The network identified by the joint analysis of common and rare variants also identified genes for both oxidative and non-oxidative ethanol metabolism, including *ADH, ALDH, UGT, CYP*, and *SULT* family genes. Ethanol is primarily metabolized in the liver, but is also metabolized by the stomach and the brain (Zakhari, 2006), which was reflected in the high enrichment of network genes in the liver, gastrointestinal tissues, and the brain. Disulfiram, by inhibiting *ALDH1A1* - which was a gene in the alcohol consumption network - is an effective treatment for alcohol use disorder (Lanz *et al*., 2023), suggesting that other genes identified by our network could also be viable pharmacological targets.

In addition to ethanol metabolism, genes found by our analyses have also been associated with neuropsychiatric conditions that are correlated (Walters *et al*., 2018) and highly comorbid with alcohol use disorder, such as depression (Ribadier and Varescon, 2019), schizophrenia (Johnson *et al*., 2023), bipolar disorder (Grunze *et al*., 2021), neuroticism (Ribadier and Varescon, 2019), and cognitive dysfunction (Nunes *et al*., 2019). For example, the rare variant analysis identified *KIF21B (Asselin et al*., *2020)*, which has been associated with smoking initiation (Saunders *et al*., 2022), ADHD (Cross-Disorder Group of the Psychiatric Genomics Consortium, 2013), and schizophrenia (Trubetskoy *et al*., 2022). *GIGYF1* has been associated with Alzheimer’s disease (Burdett *et al*., no date) and schizophrenia (Ding *et al*., 2023). Finally, *SCN7A* has been associated with unipolar depression and educational attainment (Almomani *et al*., 2023). Similarly, the alcohol consumption network identified genes that have also been associated with neuropsychiatric conditions, such as genes from the FOXP family (i.e., *FOXP1, FOXP2, FOXP4 (Sollis et al*., *2023))*. Another example is *CACNB3* and *CACNG4*, calcium channel genes that have been associated with bipolar disorder and major depression((Sklar *et al*., 2012; Marshe *et al*., 2021). Finally, the gene *ADGRG6*, which was identified by the alcohol consumption network, has been associated with depression and smoking initiation (Sollis *et al*., 2023). Integrative analyses may help clarify the shared mechanisms of these conditions, but together this emphasizes shared genetic susceptibility across these traits.

While this study found that common and rare variants that were associated with alcohol consumption identified a shared network, there are several limitations to consider. We found that *ADH1C* is needed for network colocalization, showing that it is a hub gene for this network; this highlights the need for increased power of rare variants. We only studied alcohol consumption, however future study should also consider other AUD-relevant phenotypes. Similarly, methods for mapping common SNP to genes are imperfect; we used MAGMA but other more or less stringent methods might produce different results. Additionally, we used a lenient significance threshold to select rare variants (FDR>0.25), which likely introduced some false positives into the network analysis. However, we repeated this analysis with a more stringent cutoff for rare variants (*p*_SKAT-O_<2.5×10^−7^) and found little change in significance of network overlap. Additionally, NetColoc is robust to false positives, but functions best with a moderate number of input genes (Rosenthal *et al*., 2023).

While future improvements to our methodology and the underlying data will improve our ability to understand rare and common variant interaction, this work identified the first gene network from common and rare variants of alcohol consumption.

## Materials and Methods

### Lead Contact

Further information and requests for resources should be directed to.

## Materials availability

This study did not generate new unique reagents.

## Data and code availability

All code used for analysis and data visualization is freely available in public repositories. All original code is publicly available at https://github.com/BSLeger/rare_common_alcohol_comparison.

Any additional information required to reanalyze the data reported in this paper is available from the lead contact upon request.

## Data acquisition

### Common variant experimental and control data acquisition

The common variant summary statistics for alcohol consumption were obtained from the GWAS & Sequencing Consortium of Alcohol and Nicotine use. Summary statistics were computed via a meta-analysis of GWAS results representing 666,978 individuals of European ancestry (Saunders *et al*., 2022). Summary statistics for common variant negative control traits were obtained from the Neale Lab Round 2 GWAS (Abbott *et al*., 2022) (http://www.nealelab.is/uk-biobank). Phenotype codes are FEV1: Forced Expiratory Volume per Second (20153_irnt) and Heel Quantitative Ultrasound Index (QUI) (4104_irnt). These negative control traits were selected as they have similar numbers of implicated genes to alcohol consumption (250<N<350), similar SNP heritability to alcohol consumption (*h*^*2*^ >0.30), and minimal genetic correlation with a comparable alcohol consumption trait (Amount of Alcohol Drunk on a Typical Drinking Day (20403)) (|*r*_g_| < 0.38). These estimates were obtained from the UKB Heritability browser (https://nealelab.github.io/UKBB_ldsc/h2_browser.html) and UKB Genetic Correlation browser (https://ukbb-rg.hail.is/), both generated by the Neale Lab (Abbott *et al*., 2022).

### Rare variant and gene experimental and control data acquisition and filtering

Rare variant data was downloaded from Genebass’s Hail library (gs://ukbb-exome-public/500k/results/variant_results.mt), and queried for alcohol consumption by phenotype code (alcohol_intake_custom) using Hail. To increase the confidence in rare variants, we selected for alcohol consumption rare variants that have MAC in the top 50% (MAC>2). Rare variant gene-level associations was downloaded from Genebass browser (https://app.genebass.org/). Due to limited comparisons between rare variant datasets, rare variant controls were filtered based on heritability and genetic correlations listed above, which were calculated based on common variants. Rare variant controls were chosen if they had heritability greater than 0 (*h*^*2*^ >0.01), minimal genetic correlation with a comparable alcohol consumption trait (Amount of Alcohol Drunk on a Typical Drinking Day (20403)) (0.0< |*r*_g_| < 0.2), and had the minimum number of rare seed genes recommended for network propagation using NetColoc (n>5), using comparable significance cutoffs as used for alcohol consumption rare seed genes (false discovery rate corrected burden or SKAT-O or SKAT<0.25, calculated for each individual dataset). Phenotype codes are as follows: Alcohol Consumption (alcohol_intake_custom), FEV1: forced expiratory volume per second (20153), Pulse Rate (4194), Heel bone mineral density (BMD) T-score, automated (78), Other malignant neoplasms of skin (C44), Malignant neoplasm of breast (C50). The stringent alcohol consumption rare seed genes were selected if the genes were significant by any test (*p*<0.05) after bonferroni correction (*p*_SKAT-O_<2.5×10^-7^; *p*_burden_<6.7×10^−7^, *p*_SKAT_<2.5×10^−7^). Genes were considered leniently significant if any of the gene tests identified the genes as significant (*p*<0.25) after false discovery rate correction (*p*_SKAT-O_<1.5×10^-4^; *p*_burden_<1.1×10^−4^, *p*_SKAT_<2.7×10^−5^).

### Molecular Interaction Networks

The Parsimonious Composite Network (Huang *et al*., 2018) (PCNet v1.4) was obtained from the network data exchange (NDEx, ndexbio.org), UUID: c3554b4e-8c81-11ed-a157-005056ae23aa. PCNet is a molecular interaction resource formed from integrating 21 interaction databases that contain various evidence types, including physical protein-protein, genetic, co-expression, and co-citation evidence. Each interaction in PCNet is supported by at least two of the component databases, a threshold chosen to maximize the ability of PCNet to perform gene set recovery tasks via network propagation. All seed genes were mapped to the nodes of PCNet via gene symbols.

### Common variant gene mapping

We generated gene-level significance values from the SNP-level summary statistics using the MAGMA algorithm (de Leeuw *et al*., 2015) using default parameters. Annotation windows were 10 kb, and the 1000 Genomes European reference panel was used for genome, and Hg38 Gene locations, downloaded from MAGMA’s launch page (https://ctg.cncr.nl/software/magma). MAGMA projects the SNP matrix onto principal components, and uses the principal components to predict for the phenotype using linear regression. Association of the gene to the phenotype using the principal component SNP matrix is used to calculate an F statistic, which is used to calculate the *p*-value for the individual genes. Genes were considered significant if they were *p*<2.6×10^−6^.

### Generation of the alcohol consumption network

#### Network propagation and co-localization

We used the Python package NetColoc (Rosenthal *et al*., 2023) (https://pypi.org/project/netcoloc/) for network propagation and co-localization. The sets of significant trait-associated genes from GWAS were used as “seed” genes for network propagation using a Random Walk with Restart (Vanunu *et al*., 2010) algorithm. Following network propagation with *α*= 0.5, we calculated a network proximity score (NPS) for each gene in the network by comparing the observed results to a null distribution. The null distribution was formed by propagating 1,000 randomly selected seed gene sets. Each set was sampled to preserve the size and degree distribution of the original input set. As previously implemented (Rosenthal *et al*., 2021, 2023; Wright *et al*., 2023), we binned all genes in the network by degree with a minimum of 10 nodes per bin. For each gene, the NPS was calculated as a z-score comparing the observed heat at that gene after network propagation of the gene set, to the mean of the null distribution heats at that gene. All heat values are log-transformed to ensure the distributions are approximately normal.

NetColoc recommends fewer than 500 input seed genes given the sample space of PCNet (∼18,000 genes). Therefore, we employed a weighted sampling procedure for any trait having more than 500 significantly associated genes. We sampled 500 genes from the set of all significant genes (weighted by –log_10_p from GWAS) and ran the propagation analysis from this subset. After 100 repetitions, the 75% percentile NPS score was selected to approximate a consensus score for each gene.

From input seed genes from common and rare variants, we independently calculated NPS_common_ and NPS_rare_ for each trait. We then defined a gene as colocalized between both if it had high proximity to both input sets. Therefore, we defined the combined network proximity NPS_common-rare_ as the product of the independent dataset vectors:

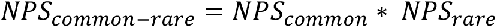

### Definition of the alcohol consumption network

From NPS_common-rare,_ we selected genes with high proximity scores from both common and rare sources to define the network using the following thresholds: NPS_common-rare_ > 3, NPS_common_ > 1.5, and NPS_rare_ > 1.5. To calculate the significance of the network co-localization, we compared the conserved network size and the mean NPS_common-rare_ to a permuted null distribution. We permuted the labels of NPS_rare_ and NPS_common_ 10,000 times, and each time calculated the mean NPS_common-rare_ across all genes and the number of genes passing the above thresholds. For genes present in both input sets, labels were permuted separately to maintain the higher expected distribution for these genes. The significance of the conserved network size and mean NPS_common-rare_ was calculated by Z-test.

### Validation and functional annotation

#### Gene family annotation

Gene families were manually assigned based on gene families identified from the Uniprot ID mapping function (https://www.uniprot.org/id-mapping) on 28 November 2023. Families were assigned broadly to make functional groups more evident, with a minimum of 2 genes per family required to be labeled.

#### GWAS catalog

To identify previously annotated genes, we used GWAS findings aggregated by the GWAS catalog (https://www.ebi.ac.uk/gwas/). The GWAS catalog’s gene level associations v.1.0.2 were downloaded on 2 August 2023. We identified genes that had previously been associated with various traits by querying the Mapped Trait and the Disease/Trait for various keywords (see github for specific parameters). Traits were grouped into alcohol use traits, smoking and nicotine use traits, non-alcohol or smoking substance use disorders (for example, opioid use disorder), and non-SUD neurological and psychiatric traits (including cognitive traits, mental health and psychiatric traits, and neuro-degenerative traits). All groups are mutually exclusive. Within each group, traits are listed as Mapped Trait: Disease/Trait for clarity, and listed only once per gene for readability. Enrichment for each group was calculated using a hypergeometric test. Genes mentioned in text were reconfirmed using the GWAS catalog browser.

#### Rare Variant PheWAS

To assess the association of network genes with other phenotypes through rare variants, gene level PheWAS results were downloaded from Genebass’s Hail database (gs://ukbb-exome-public/500k/results/results.mt). Phenotypes were mapped to network genes by gene symbol. Phenotypes were determined as significant using the same *p*-value cutoffs as used for lenient seed genes from alcohol consumption (*p*_SKAT-O_<1.5×10^−4^; *p*_burden_<1.1×10^−4^, *p*_SKAT_<2.7×10^−5^).

#### Tissue Enrichment

To assess the tissue-specific expression of network genes, we used the Functional Mapping and Annotation of Genome-Wide Association Studies (FUMA) suite’s gene to function tool (Watanabe *et al*., 2017). We used FUMA to calculate the enrichment of gene sets for 54 tissue types from human GTEx v8 (GTEx Consortium, 2020). As described previously, this method takes normalized gene expressions (reads per kilobase per million, RPKM) from each GTEx tissue, and maps these genes to entrez ID (Watanabe *et al*., 2017). Pre-calculated differentially expressed genes (DEG) sets were defined using a two-sided t-test per label versus all remaining tissue types. Genes with a Bonferroni corrected *p*-value<0.05 and absolute log fold change≥0.58 were selected as DEG. For the signed DEG, the direction of expression was taken into account. The -log 10(*p*-values) in the graph were calculated by hypergeometric test (Watanabe *et al*., 2017).

## Acknowledgments

Montana Kay Lara contributed scientific input to this manuscript. BSL was supported in part by NIGMS T32 GM008666. SSR was supported by NIH/NIDA DP1DA054394. SSR and AAP were supported by NIAAA R01AA029688.

## Author Contributions

BSL, SSR, and AAP conceptualized the study. BSL and JJM performed analysis. All authors wrote and edited the manuscript.

## Supplemental Figures

**Supplemental Figure 1.**
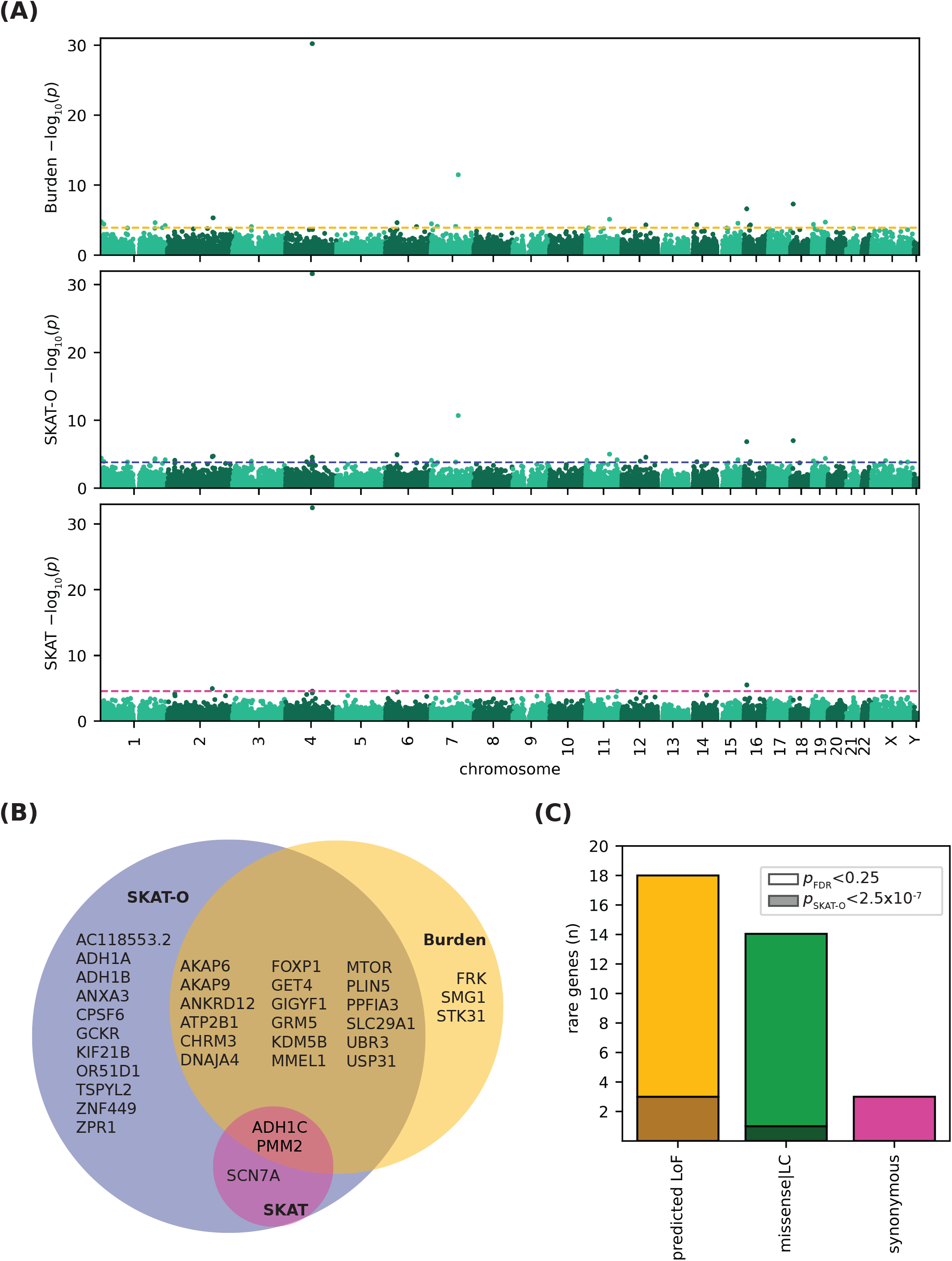
Rare variants-implicated genes mediating alcohol consumption. (**A**) Manhattan plot of association with alcohol consumption for rare-variant implicated genes calculated by burden test (top), SKAT-O (middle), and SKAT (bottom). Dotted lines indicate FDR<0.25 cutoff for each test. (**B**) Venn diagram of leniently significant (FDR<0.25) genes identified from rare variants for alcohol consumption, broken down by SNP to gene algorithm used (burden, SKAT, SKAT-O). (**C**) Stacked bar chart of rare seed gene mutation type, grouped by SNP to gene algorithm. FDR<0.25 genes are shown in light colors, and *p*_SKAT-O_<2.5x10-7 genes are shown in dark colors. No *p*_SKAT-O_<2.5x10-7 genes were annotated as synonymous.

**Supplemental Figure 2.**
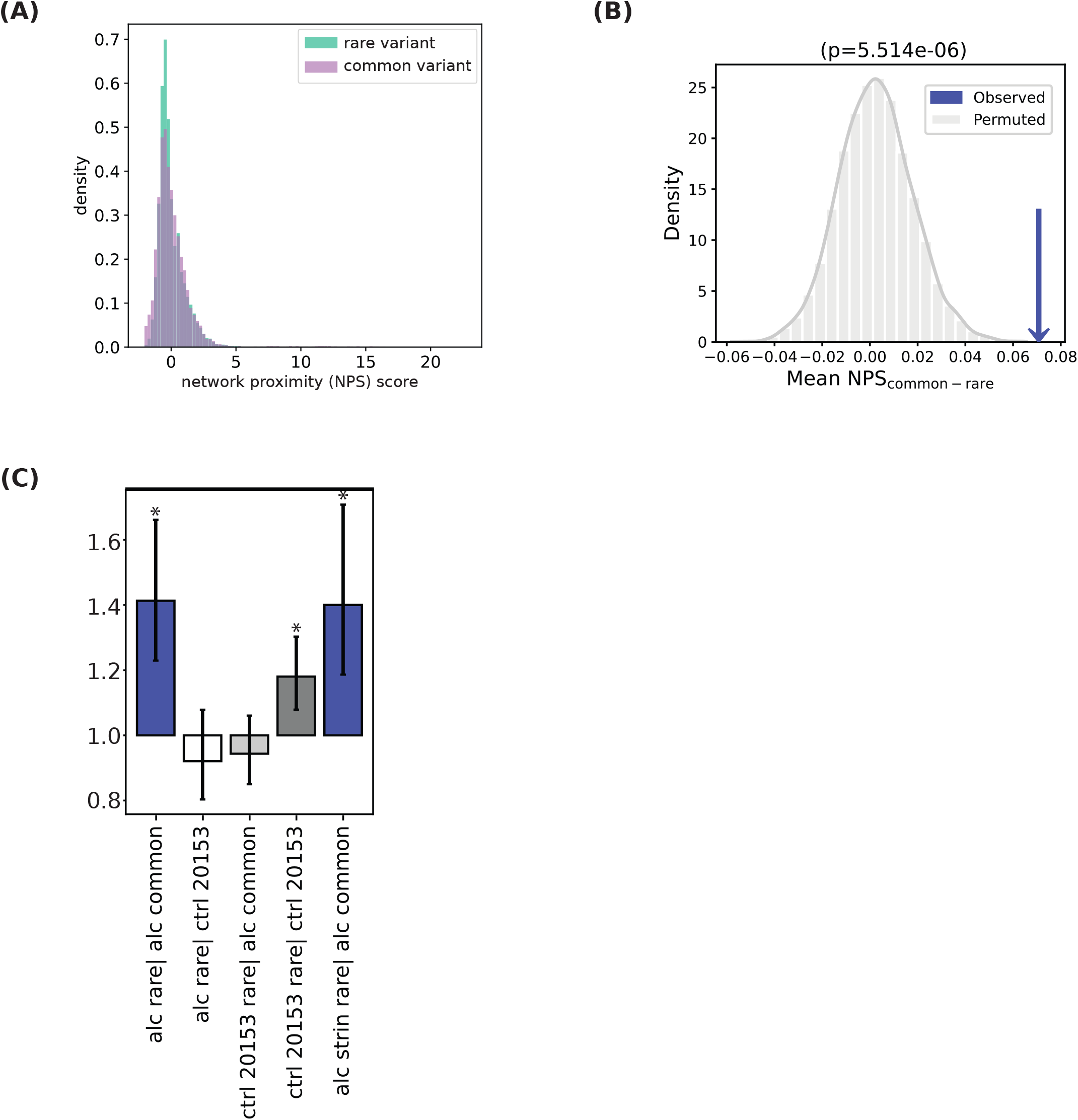
Network colocalization of rare and common alcohol consumption seed genes. (**A**) Distribution of NPS_common_ and NPS_rare_ for all nodes in PCNet. (**B**) Observed (blue arrow) and expected mean NPS_comon-rare_ for colocalization of common and rare seed genes, with significance assessed by Z-test. (**C**) The observed-to-expected ratio of colocalized network size from the following sources: alcohol consumption common and rare seed genes (left blue bar), negative control FEV1 (forced expiratory volume per second) common seed genes and alcohol consumption rare seed genes (white), FEV1 rare seed genes with alcohol consumption common seed genes (light gray), FEV1 rare and common seed genes (dark gray), and alcohol consumption stringent (*p*_SKAT-O_<2.5x10-7) rare seed genes and common seed genes. Vertical bars indicate 95% confidence intervals. Significance calculated by Z-test, Bonferroni corrected. See Table S5 for additional controls.

**Supplemental Figure 3.**
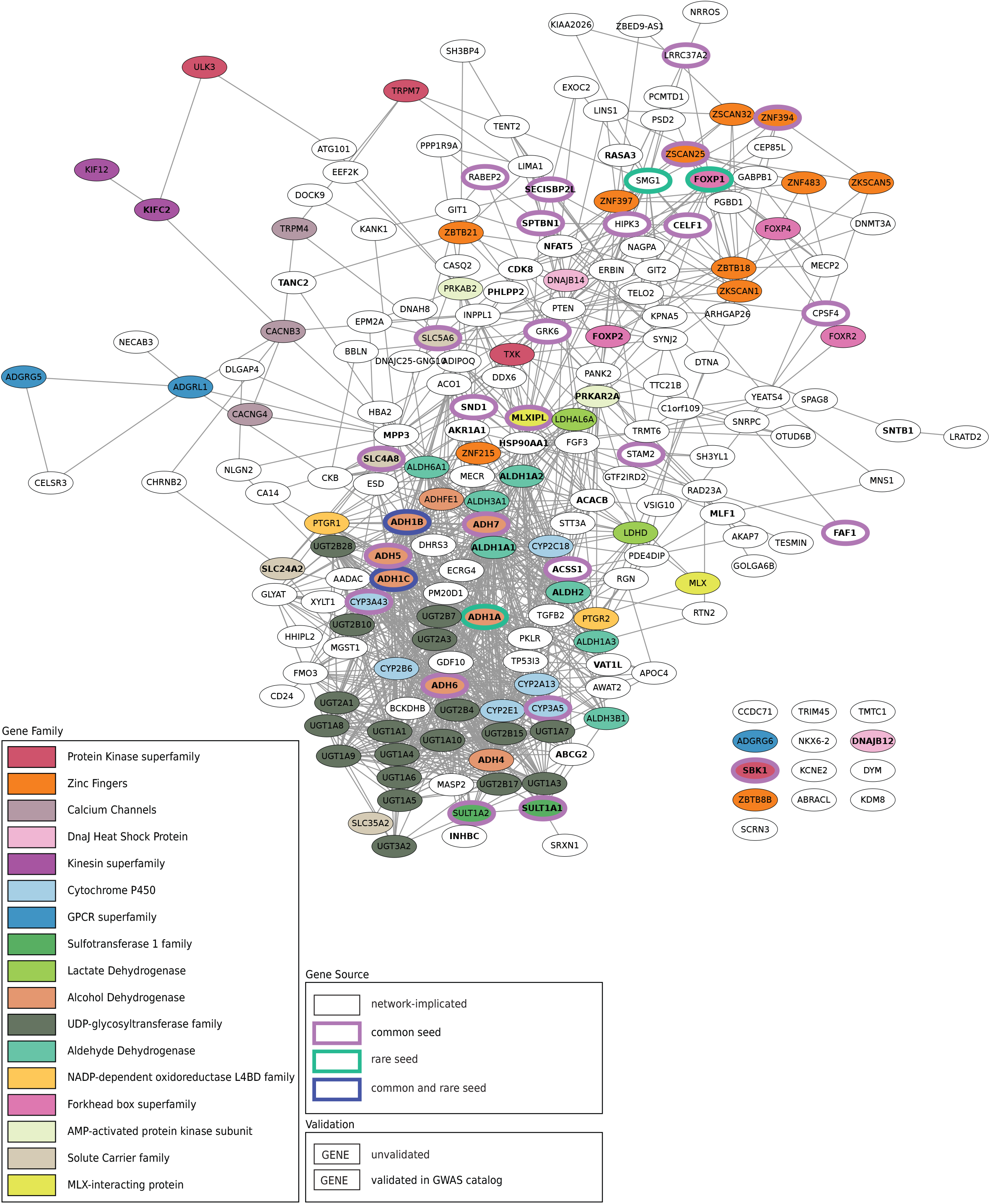
Gene families in the alcohol consumption network. Subnetwork of PCNet including all genes proximal to both common and rare seed genes, as in Figure 3. Edges maintained from PCNet. Purple outlined nodes indicate common seed genes, green outlined nodes indicate rare seed genes, dark blue outlined nodes indicate seeds from both sources. Nodes with gene symbols in bold have previously been annotated in the GWAS catalog for alcohol related phenotypes. Gene families and functional groups were manually identified, and are indicated by color, as shown in legend. Only families with 2 or more genes present in the network were annotated.

